# Ecological and genomic attributes of novel bacterial taxa that thrive in subsurface soil horizons

**DOI:** 10.1101/647651

**Authors:** Tess E. Brewer, Emma L. Aronson, Keshav Arogyaswamy, Sharon A. Billings, Jon K. Botthoff, Ashley N. Campbell, Nicholas C. Dove, Dawson Fairbanks, Rachel E. Gallery, Stephen C. Hart, Jason Kaye, Gary King, Geoffrey Logan, Kathleen A. Lohse, Mia R. Maltz, Emilio Mayorga, Caitlin O’Neill, Sarah M. Owens, Aaron Packman, Jennifer Pett-Ridge, Alain F. Plante, Daniel D. Richter, Whendee L. Silver, Wendy H. Yang, Noah Fierer

**Author notes:** Corresponding authors: Tess Brewer and Noah Fierer, University of Colorado, CIRES Building, Rm. 318, 216 UCB, Boulder, CO 80309-0216 USA.

## Abstract

While most bacterial and archaeal taxa living in surface soils remain undescribed, this problem is exacerbated in deeper soils owing to the unique oligotrophic conditions found in the subsurface. Additionally, previous studies of soil microbiomes have focused almost exclusively on surface soils, even though the microbes living in deeper soils also play critical roles in a wide range of biogeochemical processes. We examined soils collected from 20 distinct profiles across the U.S. to characterize the bacterial and archaeal communities that live in subsurface soils and to determine whether there are consistent changes in soil microbial communities with depth across a wide range of soil and environmental conditions. We found that bacterial and archaeal diversity generally decreased with depth, as did the degree of similarity of microbial communities to those found in surface horizons. We observed five phyla that consistently increased in relative abundance with depth across our soil profiles: Chloroflexi, Nitrospirae, Euryarchaeota, and candidate phyla GAL15 and Dormibacteraeota (formerly AD3). Leveraging the unusually high abundance of Dormibacteraeota at depth, we assembled genomes representative of this candidate phylum and identified traits that are likely to be beneficial in low nutrient environments, including the synthesis and storage of carbohydrates, the potential to use carbon monoxide (CO) as a supplemental energy source, and the ability to form spores. Together these attributes likely allow members of the candidate phylum Dormibacteraeota to flourish in deeper soils and provide insight into the survival and growth strategies employed by the microbes that thrive in oligotrophic soil environments.

**Importance:** Soil profiles are rarely homogeneous. Resource availability and microbial abundances typically decrease with soil depth, but microbes found in deeper horizons are still important components of terrestrial ecosystems. By studying 20 soil profiles across the U.S., we documented consistent changes in soil bacterial and archaeal communities with depth. Deeper soils harbored distinct communities compared to the more commonly studied surface horizons. Most notably, we found that the candidate phylum Dormibacteraeota (formerly AD3) was often dominant in subsurface soils, and we used genomes from uncultivated members of this group to identify why these taxa are able to thrive in such resource-limited environments. Simply digging deeper into soil can reveal a surprising amount of novel microbes with unique adaptations to oligotrophic subsurface conditions.

## Introduction

Subsurface soils often differ from surface horizons with respect to their pH, texture, moisture levels, nutrient concentrations, clay mineralogy, pore networks, redox state, and bulk densities. Globally, the top 20 cm of soil contains nearly five times more organic carbon (C) than soil in the bottom 20 cm of meter-deep profiles (1). In addition, residence times of organic C pools are typically far longer in deeper soil horizons (2), suggesting that much of the soil organic matter found in the subsurface is not readily utilized by microbes. Unsurprisingly, the strong resource gradient observed through most soil profiles is generally associated with large declines in microbial biomass (3-8); per gram soil, microbial biomass is typically one to two orders of magnitude lower in the subsurface than surface horizons (4, 6, 7). Although microbial abundances in deeper soils are relatively low on a per gram soil basis, the cumulative biomass of microbes inhabiting deeper soil horizons can be on par with that living in surface soils, owing to the large mass and volume of subsurface horizons (3, 5). Moreover, those microbes living in deeper horizons can play important roles in mediating a myriad of biogeochemical processes, including processes associated with soil C and nitrogen (N) dynamics (9, 10), soil formation (11), iron redox reactions (12, 13), and pollutant degradation (14).

Given that soil properties typically change dramatically with depth, it is not surprising that the composition of soil microbial communities also generally changes with depth through a given profile (4-6, 8, 15, 16). In some cases, the differences observed in microbial communities with depth through a single soil profile can be large enough to be evident even at the phylum level of resolution. For example, both Chloroflexi (15, 17) and Nitrospirae (15) may increase in relative abundance with depth. However, while previous work suggests that particular taxa can be relatively more abundant in deeper soils, it is unclear if such patterns are consistent across distinct soil and ecosystem types. We hypothesized that there are specific groups of soil bacteria and archaea that are typically rare in surface horizons, but more abundant in deeper soils. Taxa that are proportionally more abundant in deeper soil horizons likely have slow-growing, oligotrophic life history strategies due to the lack of disturbance at depth and the low resource conditions typical of most deeper soil horizons (18). Likewise, we expect deeper soils to harbor higher proportions of novel and undescribed microbial lineages given that oligotrophic taxa are typically less amenable to *in vitro*, cultivation-based investigations (19).

We designed a comprehensive study to investigate how soil bacterial and archaeal communities change with soil profile depth, to identify taxa that are consistently more abundant in deeper horizons, and to determine what life history strategies enable these taxa to thrive in the resource-limited conditions typical of most subsurface horizons. We collected soil samples at 10-cm increments from 20 soil profiles representing a wide range of ecosystem types throughout the U.S., with most of the profiles sampled to one meter in depth. We examined the bacterial and archaeal communities of these soil profiles by pairing amplicon 16S rRNA gene sequencing with shotgun metagenomic sequencing on a subset of samples. We found that deeper soil horizons typically harbored more undescribed bacterial and archaeal lineages, and we identified specific phyla (including Dormibacteraeota, GAL15, Chloroflexi, Euryarchaeota, and Nitrospirae) that consistently increased in relative abundance with depth across multiple profiles. Moreover, we found one candidate phylum (Dormibacteraeota, formerly AD3) to be particularly abundant in deeper soil horizons with low organic C concentrations. From our metagenomic data, we were able to assemble genomes from representative members of this candidate phylum and document the life history strategies, including low maximum growth rates and spore-forming potential, that are likely advantageous under low resource conditions.

## Results & Discussion

### Sample descriptions and soil properties linked to soil depth

We collected soils from a network of 10 current and former Critical Zone Observatories (CZOs) located across the U.S. (Figure 1a) that span a broad range of hydrogeological provinces, soil orders, and ecosystem types, including tropical forest, temperate forest, grassland, and cropland sites. Soils were sampled from two distinct profiles per CZO for a total of 20 different soil profiles. Details of the site characteristics and edaphic properties for each of the 20 soil profiles are provided in Supplemental Dataset 1. Soils were collected from the first meter (where possible) of freshly excavated profiles, sampling at 10 cm increments and focusing on mineral soil horizons only (O horizons, if present, were not sampled). Together, this collection yielded 179 individual soil samples collected across sites with a wide range of different climatic conditions (e.g., mean annual temperatures ranging between 5 - 23 °C and mean annual precipitation ranging from 26 - 402 cm y^-1^, Supplemental Dataset 1). The sampled profiles ranged from poorly developed Entisols and Inceptisols to highly developed Oxisols and Ultisols (as per the U.S. Soil Taxonomy system), and reflected an extremely broad range of soil properties. For example, in the 0-10 cm depth increment, soil pH ranged from 3.3 to 9.8, organic carbon concentrations ranged from 1.3% to 21.6%, and texture ranged from 0% to 45% silt + clay across the profiles.

**Figure 1:**
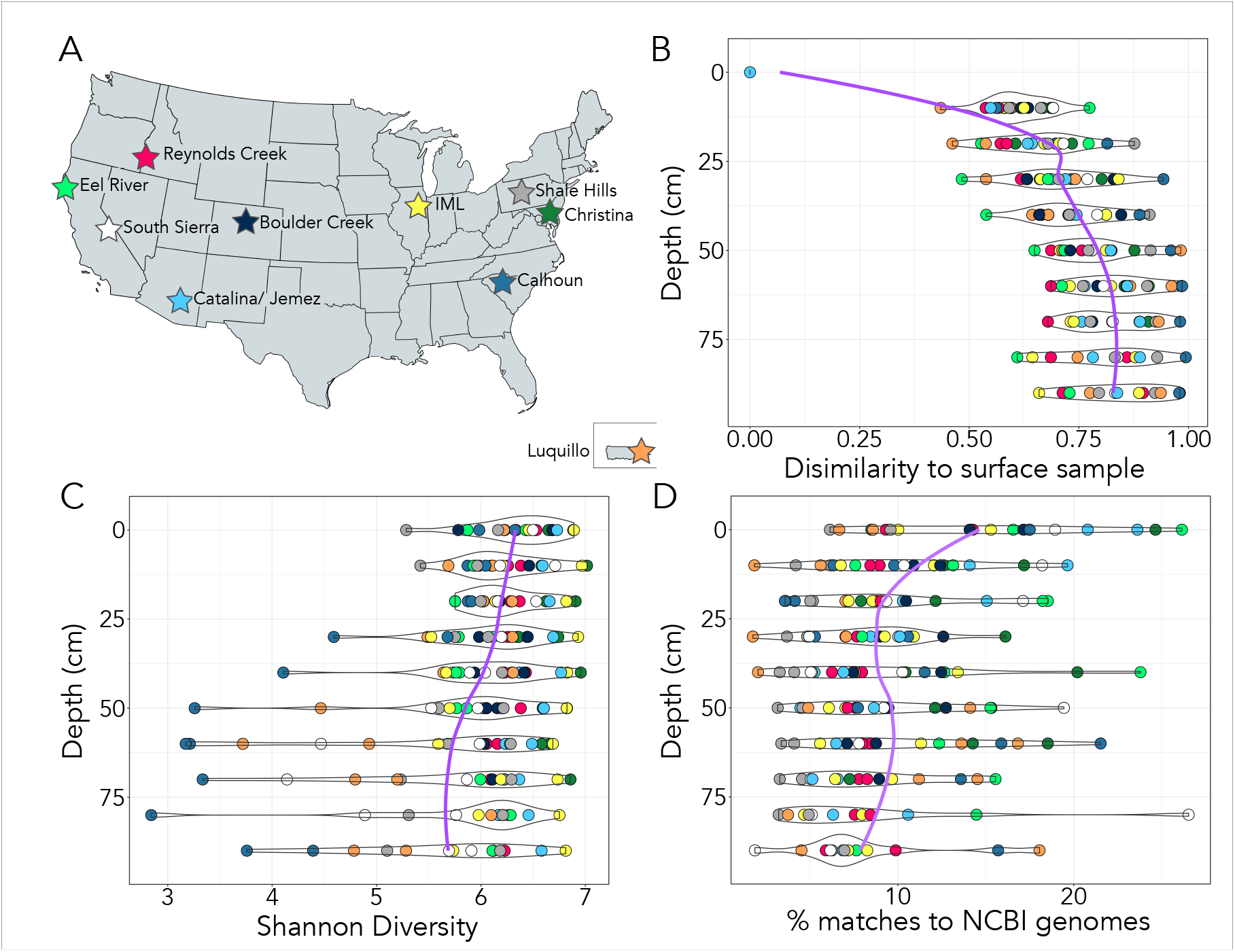
(A) Site map of sampling locations. We analyzed bacterial and archaeal communities from 2 soil profiles located at each of 10 different CZOs across the U.S. Each profile was sampled in 10 cm intervals from surface soils to one meter in depth (where possible). **(B) Bray-Curtis dissimilarity to surface samples increases with depth**. As depth increases, soil bacterial and archaeal communities become less similar to those communities at the surface. **(C) Bacterial and archaeal diversity generally decreases with depth.** Colors of points match the colors of the CZO sites indicated in panel A with two profiles sampled per site (N=20). **(D) The proportion of 16S rRNA gene sequences from the sampled soils for which representative genome data are available decreases with depth.** We matched our 16S rRNA gene amplicon sequences to 16S rRNA genes from finished bacterial and archaeal genomes in the NCBI database. At deeper soil depths, we found that fewer taxa in our dataset had representative genomes, indicating that the bacterial and archaeal taxa found in deeper soil horizons are less represented in genomic databases than those found in surface soils.

Some soil properties changed consistently with depth across all 20 profiles. Total N and organic C concentrations were both negatively correlated with soil depth, in agreement with previous observations (1, 20) (depth vs. %C rho=-0.61, p<0.001; depth vs. %N rho=-0.56, p<0.001; Spearman). On average, soil total organic C concentrations below 50 cm were 4.4 times lower than in surface soils, while total N concentrations were 6.3 times lower. While we measured a suite of additional chemical and soil properties (Supplemental Dataset 1), only clay concentrations exhibited consistent changes with depth (with percent clay generally increasing with depth; rho=0.29, p<0.001; Spearman). Given that our sampling effort included a wide range of different soil types and the expectedly high degree of variability in inter- and intra-profile edaphic characteristics, our goal was not to determine if distinct soil samples harbored distinct microbial communities or to characterize the factors related to shifts in overall community composition. Rather, our goal was to determine if there were any consistent changes in soil microbial communities with depth across the 20 sampled profiles.

### Community characteristics linked to soil depth

Unsurprisingly, we found that the location of each soil profile had a strong influence on the composition of soil bacterial and archaeal communities as determined by 16S rRNA gene amplicon sequencing (r = 0.47, p < 0.001, Permanova). Individual soil profiles generally harbored distinct microbial communities (Figure 2, Supplemental Figure 1). In addition to this variation across the profiles, soil depth also had a significant effect on the composition of the bacterial and archaeal communities within individual profiles (p < 0.01 for 16 of 20 profiles, rho values ranging from 0.24 - 0.45). In general, the variation in community composition with depth within a given profile, while significant, was less than the differences in soil communities observed across different profiles when all profiles and soil depths were examined together (Depth: r = 0.02, p < 0.001, Location: r = 0.47, p < 0.001, Permanova).

**Figure 2:**
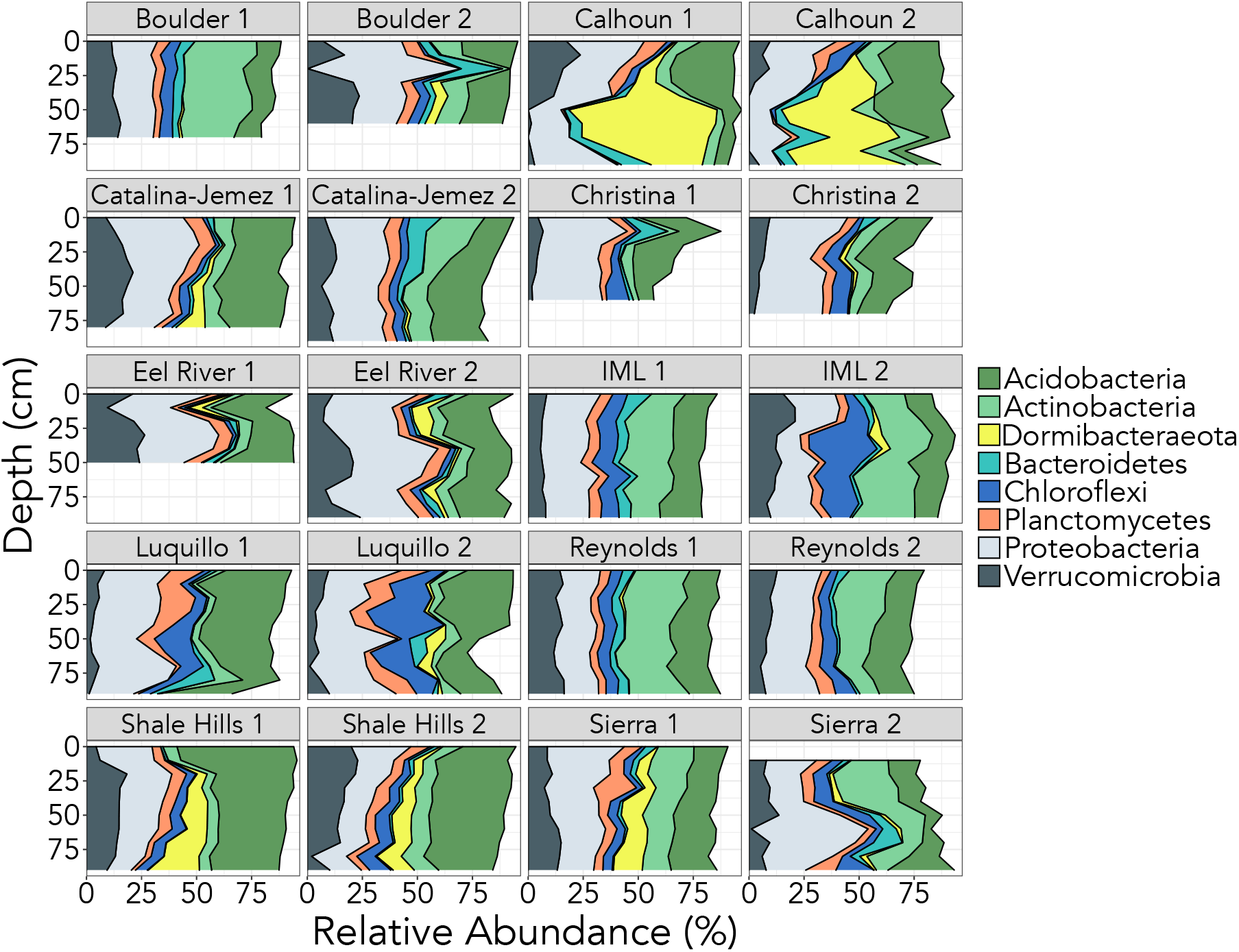
Different soil profiles have distinct microbial communities. Here we show the relative abundances of the eight most abundant phyla identified from our 16S rRNA gene amplicon sequencing effort. Not all profiles were sampled to one meter due to variable bedrock depth. Note that the two profiles sampled from each CZO site were selected to represent distinct soil types (details on soil characteristics are available in Supplemental Dataset 1).

Several characteristics of the bacterial and archaeal communities changed consistently with depth despite the high degree of heterogeneity observed across the different soil profiles. As soil depth increased, microbial communities found at depth became increasingly dissimilar to those found in surface horizons (Figure 1B). When we analyzed the entire sample set together, dissimilarity to surface soils (0-10cm depth) was positively correlated with depth (p < 0.001, rho = 0.73, Spearman). This trend also held for 17 out of 20 individual soil profiles (depth was not significant in both Eel sites and IML site 1). We also found that, in general, the diversity of microbial communities decreased with depth, with several CZOs exhibiting stronger declines with depth than others (Calhoun, Luquillo, and Southern Sierra, Figure 1C). Lastly, when we compared the 16S rRNA gene sequences from this study to those 16S rRNA gene sequences from finished bacterial and archaeal genomes in the NCBI database, we found that the proportion of taxa for which genomic data is available declined with depth (from 6.2 - 26.1% in surface soils, to 1.9 - 18.0% in the deepest horizons sampled, Figure 1D). Although representative genomes are unavailable for the majority of soil bacterial and archaeal taxa (21), genomic information from closely-related taxa is available for a smaller proportion of taxa living at depth than those found in surface soil horizons.

### Taxonomic shifts with soil depth

Although each soil profile harbored distinct microbial communities (Figure 2), we identified five phyla that consistently increased in abundance with soil depth as measured by Spearman correlations across the entire dataset: Chloroflexi, Euryarchaeota, Nitrospirae, and the candidate phyla Dormibacteraeota and GAL15 (Figure 3). For example, Dormibacteraeota were 27 times more abundant in soils at 90 cm than in surface horizons. The candidate phylum Dormibacteraeota, Chloroflexi, and Nitrospirae have previously been found to increase in abundance with increasing soil depth in individual profiles (15, 17), while candidate phylum GAL15 has been shown to be abundant in oxic subsurface sediments (22). Members of these phyla are likely oligotrophic taxa adapted to survive in the resource-limited conditions found in deeper horizons. Indeed, soil Euryarchaeota (23), Chloroflexi, and Nitrospirae (24) have been shown to decrease in abundance upon soil fertilization. These five phyla are also underrepresented in public genome databases; together, they account for only 2.8% of bacterial and archaeal genomes deposited in IMG (as of Dec 2018), reinforcing the observation highlighted in Figure 1D that poorly described taxa tend to be relatively more abundant in deeper soil horizons.

**Figure 3:**
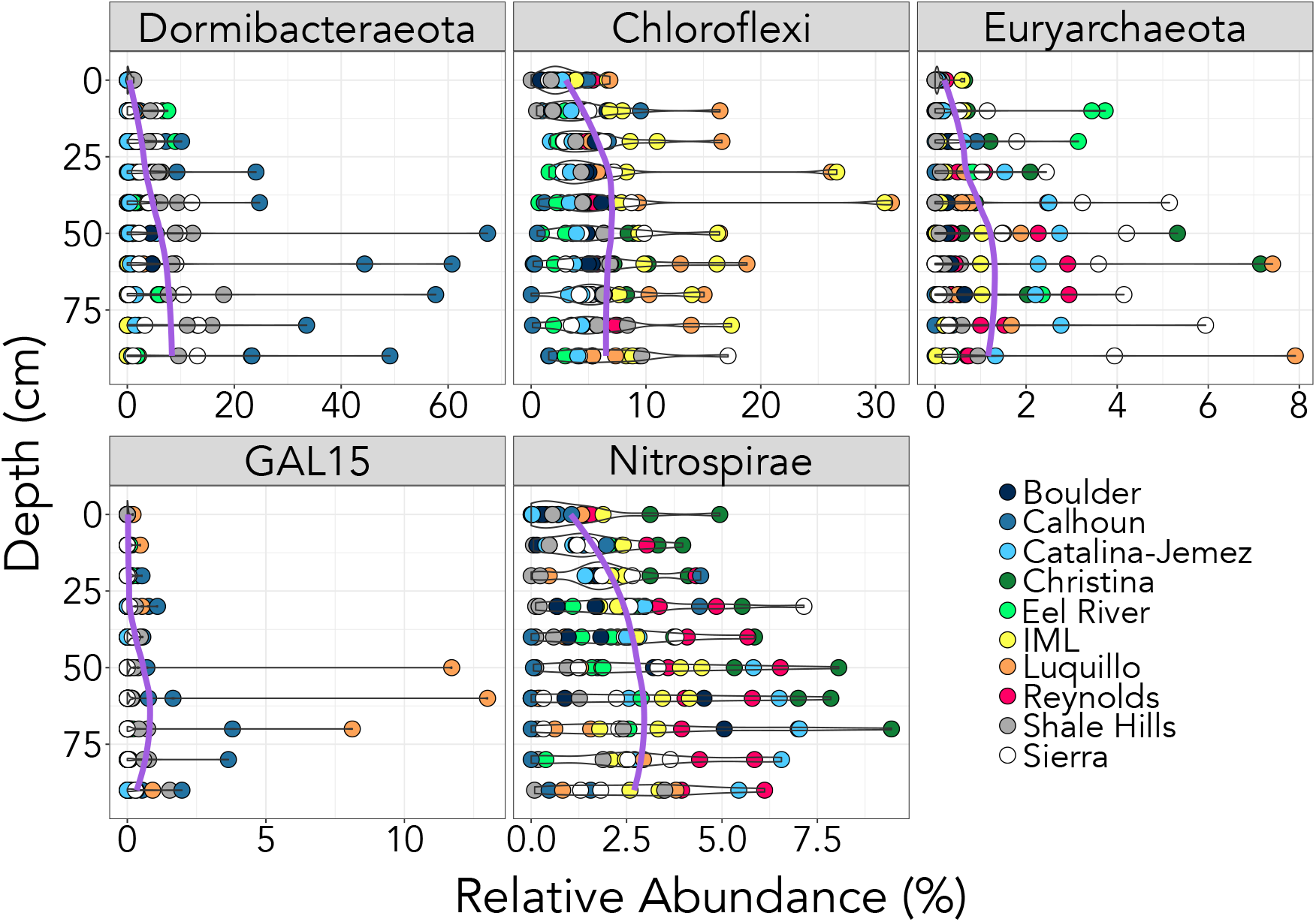
Five bacterial and archaeal phyla that consistently increased in relative abundance with soil depth. These phyla were identified via Spearman rank correlations against depth (FDR corrected p values < 0.02, rho > 0.22). For details on all phylum level abundances in each individual soil profile, see Supplementary Table 5.

### Community-level shotgun metagenomic analyses

We selected one soil profile from nine of the CZO sites for shotgun metagenomic sequencing, targeting those profiles that displayed the most dissimilarity among different depths. Together, we obtained shotgun metagenomic data from 67 soil samples with an average of 7.84 million quality-filtered reads per sample. We first used these metagenomic data to quantify changes in the relative abundances of the bacterial, archaeal, and eukaryotic domains with depth. The overwhelming majority of rRNA gene sequences that we detected were from bacteria (89.2% - 98.7% of reads), followed by archaea (0.03% - 7.70%), and then eukaryotes (0.04% - 4.27%). Interestingly, we found that the proportion of eukaryotic sequences in our samples decreased with depth (rho = -0.32, p = 0.05). Most of these eukaryotic rRNA gene reads were classified as Fungi (58%), then Charophyta (16%), Metazoa (9.3%), and Cercozoa (7.0%). These results are in line with previous work showing that the contributions of eukaryotes, most notably fungi, to microbial biomass pools typically decrease with soil depth (25).

We also directly compared the results obtained from our 16S rRNA amplicon and shotgun metagenomic sequencing across the same set of samples. We did this to check whether our PCR primers introduced significant biases in the estimation of taxon relative abundances. We found that the shotgun and amplicon-based estimations of the abundances of each of the eight phyla that were the most ubiquitous and abundant across the sampled profiles (Figure 2) were well correlated (Supplemental Figure 2, mean rho values = 0.70). Next, we checked whether our primers missed any major groups of bacteria or archaea, as it has been noted that many taxa from the Candidate Phyla Radiation (CPR, recently assigned to the superphylum Patescibacteria, (26) are not detectable with the primer set used here (27). While we found that our primer pair did fail to recover sequences from the superphylum Patescibacteria, these taxa were rare in our data - the entire superphylum accounted for only 0.5% of 16S rRNA gene reads across the whole metagenomic dataset.

### Candidate phylum Dormibacteraeota is relatively more abundant in soils with low organic carbon

We found that members of phylum Dormibacteraeota were consistently more abundant in deeper soil horizons and particularly abundant in subsurface horizons from the Calhoun and Shale Hills CZOs (Figure 4). In these soils, Dormibacteraeota dominated the microbial communities – in some samples, over 60% of 16S rRNA sequences were classified as belonging to members of the Dormibacteraeota candidate phylum. The high abundances of Dormibacteraeota were confirmed with shotgun metagenomic analyses (Supplemental Figure 2), indicating the abundances of this phylum were not inflated by PCR primer biases. The candidate phylum Dormibacteraeota was first observed in a sandy, highly weathered soil from Virginia, U.S. (28) and does not yet have a representative cultured isolate. The phylum was previously known as ‘AD3’, but was renamed Dormibacteraeota after three genomes from the phylum were assembled from Antarctic soils (29). Other representative genomes from this phylum have also become available with the recent addition of 47 genomes assembled from thawing permafrost (30). The phylum Dormibacteraeota has been observed in subsurface soil horizons previously (31, 32), and its relative abundance has been found to be negatively correlated with water content, C, N, and total potential enzyme activities (17).

**Figure 4:**
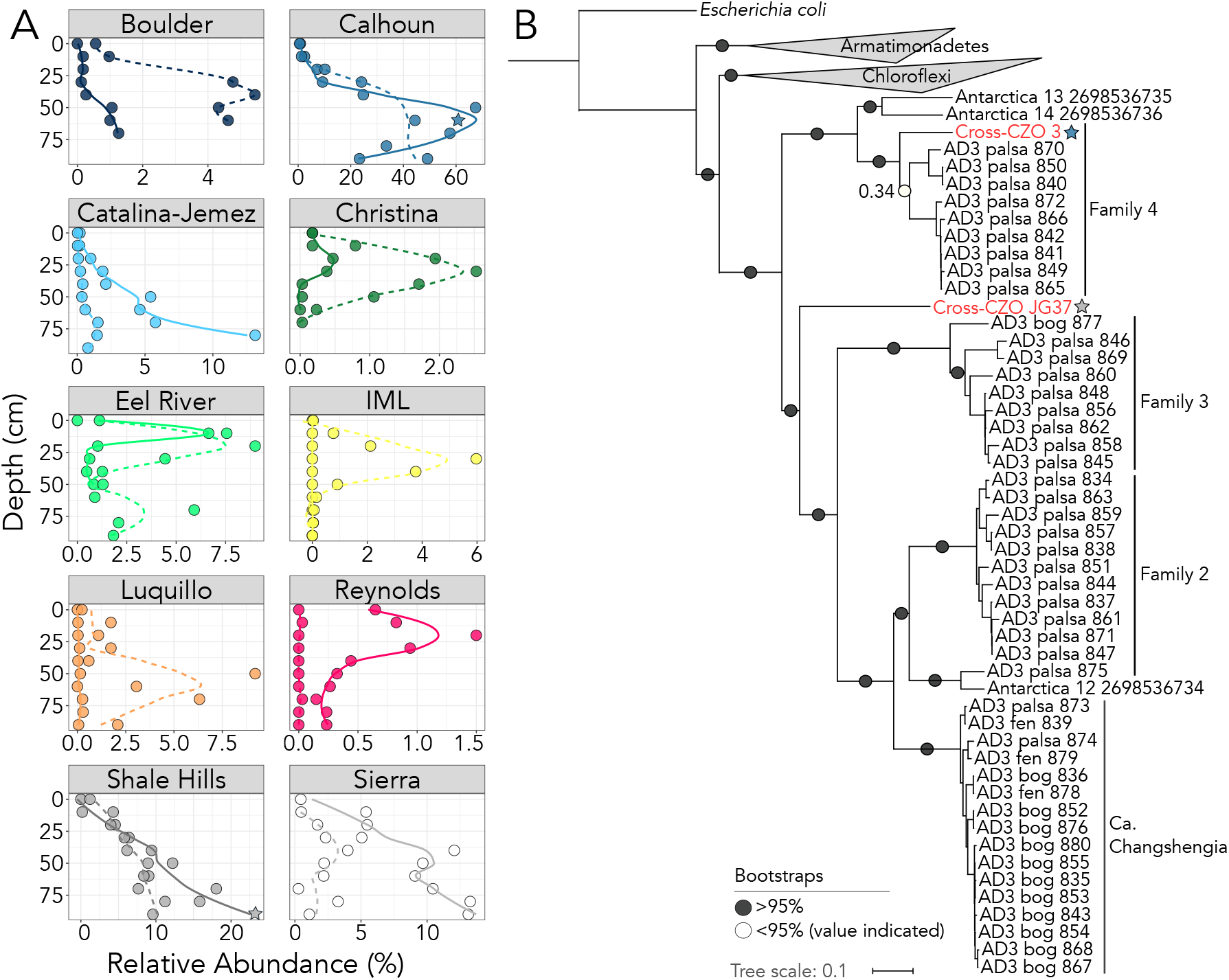
(A) The 16S rRNA gene relative abundance of phylum Dormibacteraeota is variable across different soil profiles, but generally increases with depth. The samples used for the Dormibacteraeota genome assemblies are noted with stars. **(B) The two Dormibacteraeota genomes we assembled from soil profile metagenomic data cluster phylogenetically with previously published Dormibacteraeota genomes.** Our deep soil genomes also fall near the known sister phyla Chloroflexi and Armatimonadetes, validating their identity as members of candidate phylum Dormibacteraeota. This tree was created using the concatenated marker gene phylogeny generated from GTDBTk (26), and was plotted using ITOL (64). Only closely related phyla are included in the tree. Genomes assembled in this study are indicated in red and all other AD3/Dormibacteraeota genomes originated from either (29) or (30). The family groupings for the Dormibacteraeota tree were first presented in (30).

While the abundance of phylum Dormibacteraeota was generally positively correlated with depth across all samples included in this study (rho = 0.22, p = 0.02, Spearman), this pattern did not hold for all profiles (Figure 4). Instead, we found organic C concentrations to be the best predictor of the abundance of Dormibacteraeota in these soil communities (Supplemental Figure 3); Dormibacteraeota were typically eight times more abundant in soils with less than 1% organic C than in soils where organic C concentrations were greater than 2%. Because soil depth and organic C concentrations were correlated across the profiles studied here, we used an independent dataset of surface soils (0-10 cm) collected from 1006 sites across Australia to determine if the abundances of Dormibacteraeota were also correlated with organic C concentrations when analyses were restricted to a broad range of distinct surface soils (33). Indeed, we found that the relative abundances of Dormibacteraeota in Australian surface soils (which ranged from 0.0 to 7.0% of 16S rRNA gene sequences) were also negatively correlated with soil organic carbon concentrations (Supplemental Figure 3). Together these results indicate that members of the Dormibacteraeota phylum are typically most abundant in surface or subsurface soils where organic C concentrations are relatively low.

### Dormibacteraeota draft genomes recovered from metagenomic data

To gain more insight into the potential traits and genomic attributes of soil Dormibacteraeota, we conducted deeper shotgun metagenomic sequencing on several soils where Dormibacteraeota were found to be particularly abundant (Figure 4A) with the goal of assembling draft genomes from members of this group. We assembled two Dormibacteraeota genomes, both from deep soils (Figure 4). These genomes are considered medium-quality drafts according to published genome reporting standards for MAGs (34); bin 3 is estimated to be 72.6% complete at 3.43 Mb, while bin JG-37 is 69.9% complete at 2.48 Mb (further genome details in Supplemental Table 1). These genomes are similar in size to those previously assembled from the phylum (range 3.0-5.3 Mb all >90% complete (29), range 1.6-4.3 Mb all > 70% complete (30)). These genomes share only 45.1% average amino acid identity (AAI) (35) and cluster phylogenetically with the Dormibacteraeota genomes assembled from Antarctic soil metagenomes (29) and those from permafrost metagenomes (30) (Figure 4B).

Analyses of the Dormibacteraeota genomes we recovered indicate that members of this phylum are aerobic heterotrophs adapted to nutrient poor conditions. Both Dormibacteraeota genomes encode high-affinity terminal oxidases, indicative of an aerobic metabolism (cbb_3_ binJG37, bd bin3). These genomes contain no markers of an autotrophic metabolism, with no RuBisCO or hydrogenase genes detected in either of the assembled genomes. Both Dormibacteraeota genomes contain trehalose 6-phosphate synthase, a key gene in the pathway for the synthesis of trehalose, a C storage compound that also confers resistance to osmotic stress and heat shock (36) and protects cells from oxidative damage, freezing, thermal injury, or desiccation stress (37). Additionally, both genomes contain glycogen catalysis (alpha-amylase, glucoamylases) and synthesis (glycogen synthase) genes. The ability to synthesize, store, and break down glycogen has been shown to promote the survival of bacteria during periods of starvation (36, 38). These attributes likely confer an advantage in resource-limited soils, as the ability to store C for later use may be advantageous in environments where organic C is infrequently available or of low quality.

Based on several lines of evidence, soil-dwelling Dormibacteraeota appear to be oligotrophic taxa with low maximum growth rates. First, as mentioned above, these taxa have the highest relative abundances in soils with low organic C concentrations where we would expect oligotrophic lifestyles to be advantageous. Second, both Dormibacteraeota genomes appear to encode a single rRNA operon, a feature often linked to low maximum potential growth rates (39). Third, although we cannot directly measure the maximum growth rate of uncultivated bacterial cells, we can estimate maximum growth rate from genomes by measuring codon usage bias with the ΔENC’ metric (40). ΔENC’ is a measure of codon bias in highly expressed genes, and has been shown to correlate strongly with growth rate for both bacteria and archaea (41). We calculated ΔENC’ for our Dormibacteraeota genomes, the Antarctic Dormibacteraeota genomes (29), the thawing permafrost Dormibacteraeota genomes (30), and a set of bacterial and archaeal genomes which matched the 16S rRNA gene amplicon sequences recovered from our soil profile samples at ≥99% sequence similarity. The ΔENC’ values for all the Dormibacteraeota genomes clustered together towards the lower end of the spectrum for our set of soil bacteria and archaea, indicating that members of the phylum Dormibacteraeota are likely to exhibit low potential growth rates (Supplemental Figure 4).

To our knowledge, all previous Dormibacteraeota genomes were recovered from either Antarctic desert (29) or permafrost soils (30), while our genomes hail from subsurface soils collected from temperate regions. Despite these disparate origins, some central characteristics of the phylum Dormibacteraeota appear to be consistent. Similar to the Antarctic Dormibacteraeota genomes, our Dormibacteraeota genomes also contained carbon-monoxide (CO) dehydrogenase genes. There are two forms of CO dehydrogenases, which differ in their ability to oxidize CO and the rate at which they do so (42). While the active site of form I is specific to CO dehydrogenases, form II active sites also occur in many molybdenum hydroxylases that do not accept CO as a substrate (42). Using sequence data from our assembled Dormibacteraeota genomes, the Antarctic Dormibacteraeota genomes, and selected CO dehydrogenase large subunit sequences (coxL), we generated a phylogenetic tree based on the amino acid sequence of coxL (Supplemental Figure 5). With these analyses, we found that both of the Dormibacteraeota genomes recovered here possess form II CO dehydrogenases, as do two of the Antarctic Dormibacteraeota genomes. Although it has been shown that form II CO dehydrogenases can permit growth with CO as a sole C and energy source in some cases (43), further work is needed to determine whether these genes allow Dormibacteraeota to actively oxidize CO or if these genes code for molybdenum-containing hydroxylases responsible for other metabolic processes (44). Interestingly, one Antarctic Dormibacteraeota genome also encodes a form I coxL, indicating that some members of this phylum are capable of CO oxidation (Supplemental Figure 5). CO oxidation with form I CO dehydrogenases has been shown to improve survival of bacterial cells in nutrient limited conditions (45).

Analyses of our assembled Dormibacteraeota genomes also reveal that these soil bacteria may be capable of spore formation. Altogether, our Dormibacteraeota genomes contain 34 spore-related genes scattered across a variety of spore generation phases (Supplemental Table 2). Nutrient limiting conditions are known to trigger spore formation (46), and sporulation can allow bacterial cells to persist until environmental conditions become more favorable. Additionally, members of the Chloroflexi, a sister phylum to Dormibacteraeota, are capable of spore formation (47). Because there are no Dormibacteraeota isolates available to test for sporulation, we adapted a method previously used in stool samples (48) to identify spore-forming taxa using a culture-independent approach. We incubated three soil samples from our study in 70% ethanol to kill vegetative cells, and then used propidium monoazide (PMA) to block the amplification of DNA from these dead cells (49). We then sequenced these soils using our standard 16S rRNA gene amplicon method both with and without the ethanol and PMA treatment. We found that the abundances of the two dominant Dormibacteraeota phylotypes were significantly higher in the spore-selected treatment than the untreated controls (Supplemental Table 3). Other known spore formers were enriched in the spore selection treatment as well, including taxa from the orders Actinomycetales, Bacillales (48), Myxococcales (50), and Thermogemmatisporales (51). While the enrichment of Dormibacteraeota in ethanol treated samples shows that these cells are hardy, it is not conclusive proof of spore formation and further testing is needed to verify our findings (there are other methods of ethanol resistance in bacteria, such as biofilm formation (52) and residence inside other cells (53)).

### Conclusions

Our results indicate that, as soil depth increases, not only do bacterial and archaeal communities become less diverse and change in composition, but novel, understudied taxa become proportionally more abundant in deeper soil horizons. We identified five poorly studied bacterial and archaeal phyla that become more abundant in deeper soils across a broad range of locations, and investigated one of these further (the candidate phylum Dormibacteraeota, formerly AD3) to determine what characteristics may allow Dormibacteraeota to survive and dominate in resource-limited soil environments. We found that members of Dormibacteraeota are likely slow-growing aerobic heterotrophs capable of persisting in low resource conditions by putatively storing and processing glycogen and trehalose. Members of this candidate phylum also contain type I and II carbon monoxide dehydrogenases, which can potentially enable the use of trace amounts of CO as a supplemental energy source. We also found that soil-dwelling Dormibacteraeota are likely capable of sporulation, another trait that may allow cells to persist during periods of limited resource availability. More generally, analyses of these novel members of understudied phyla suggest life history strategies and traits that may be employed by oligotrophic microbes to thrive under resource-limited soil conditions.

## Materials & Methods

### Sample collection and processing

Samples were collected from the network of 10 Critical Zone Observatories (CZOs, http://criticalzone.org) across the US: Southern Sierra (CA), Boulder Creek (CO), Reynolds (ID), Shale Hills (PA), Calhoun (SC), Luquillo (PR), Intensively Managed Landscapes (IL/IA/MN), Catalina/Jemez (AZ/NM), Eel River (CA), and Christina River (DE/PA). Volunteers from each CZO excavated two separate soil profiles (“sites”) selected to represent distinct soil types and landscape positions. Soils were collected at peak greenness (as estimated from NASA’s MODIS: MODerate-resolution Imaging Spectroradiometer) between April 2016 and November 2016, with the exception of the Eel River CZO samples, which were collected in May 2017. Volunteers were asked to sample in 10-cm increments to a depth of at least 100 cm or to refusal. Site details are available in Supplemental Dataset 1.

All soil samples were sent to the University of California, Riverside for processing. A portion of each field sample was sieved (< 2 mm, ASTM No. 10), homogenized, and divided into subsamples for further analyses, with subsamples stored at either 4°C, -20°C, or -80°C. For some soils (particularly some wet, finely textured depth intervals), sieving was not practical. These samples were homogenized by mixing, with larger root and rock fragments removed by hand. In addition, as samples from Shale Hills site 2 (70—100 cm depth) consisted almost entirely of medium-sized rocks, soil was collected by manually crushing rocks with a ceramic mortar and pestle; this material was then passed through a 2-mm sieve.

DNA was extracted from subsamples frozen at -20°C using the DNeasy PowerLyzer PowerSoil kit (Qiagen, Germantown, MD, USA), according to the manufacturer’s instructions with minor modifications to increase yield and final DNA concentration based on the assumption that some sites and depths would have a relatively low microbial biomass. Specifically, 0.25 g of soil was weighed in triplicate (i.e., three 0.25 g aliquots = 0.75 g total soil per sample) from one frozen aliquot of sieved soil. Extractions on each 0.25 replicate aliquot proceeded in parallel, until the stage when DNA was eluted onto the spin filter; replicates were pooled at this point onto a single filter, and extractions proceeded from this point as a single sample. In addition, the final step of elution of the DNA from the filter was conducted with 50 *µ*L of elution buffer, instead of 100 *µ*L; the initial flow-through was reapplied to the filter once to increase yield.

### Soil characteristics

Frozen subsamples (stored at -20°C) were shipped to the University of Illinois at Urbana-Champaign for characterization of soil physicochemical properties. Soil C and N concentrations were measured on freeze-dried, sieved, and ground subsamples using a Vario Micro Cube elemental analyzer (Elementar, Hanau, Germany). Approximately 1 g of each subsample was also extracted in 30 mL of 0.5 N HCl for determination of Fe(III) and Fe(II) concentrations using a modified ferrozine assay (54). Soil texture was measured on oven-dried and sieved soil following Gee and Bauder (1986).

Soil pH and gravimetric water content were measured using modified Long Term Ecological Research (LTER) protocols, as per Robertson et al. (1999). Soil pH was determined using 15 g of field-wet soil and 15 mL of Milli-Q water (Millipore Sigma, Burlington, Massachusetts), and was measured on a Hannah Instruments (Woonsocket, RI) HI 3220 pH meter with a HI 1053B pH electrode, designed for use with semi-solids. For determining gravimetric water content, we oven-dried 7 g of soil at 105 ° C, for a minimum of 24 hours.

### Amplicon-based 16S rRNA gene analyses

To characterize the bacterial and archaeal communities in each sample, we used the barcoded primer pair 515f/806r for sequencing the V4-V5 region of the 16S rRNA gene following methods described previously (23). We amplified this gene region three times per sample, combined these products, and normalized the concentration of each sample to 25 ng using SequalPrep Normalization Plate Kits (Thermo Fisher Scientific, Waltham, MA). All samples were then pooled and sequenced on the Illumina MiSeq (2×150 paired end chemistry) at the University of Colorado Next-Generation Sequencing Facility. The sample pool included several kit controls and no template controls to check for possible contamination.

Sequences were processed using a combination of QIIME and USEARCH commands to demultiplex, quality-filter, remove singletons, and merge paired end reads. Sequences were classified into exact sequence variants (ESVs) using UNOISE2 (55) with default settings and taxonomy was assigned against the Greengenes 13_8 database (56) using the RDP classifier (57). ESVs with greater than 1% average abundance across all sequenced controls were classified as contaminants and removed from further analyses, along with ESVs identified as mitochondria and chloroplast. The entire dataset was then rarefied to 3400 sequences per sample.

### Shotgun metagenomic analyses

One soil profile from each CZO was selected for shotgun sequencing - we chose the sites that exhibited the most dissimilarity in microbial community composition through the soil profile, as we were interested in changes most associated with soil depth. Using the same DNA as used for the amplicon sequencing effort, we generated metagenomic libraries using the TruSeq DNA LT library preparation kit (Illumina, San Diego, CA). All samples were pooled and sequenced on an Illumina NextSeq run using 2×150bp paired end chemistry at the University of Colorado Next-Generation Sequencing Facility. Prior to downstream analysis, we merged and quality filtered the paired-end metagenomic reads with USEARCH. After quality filtering we had an average of 8.8 million quality-filtered reads per sample (range = 1.9 -15.4 million reads, we only included samples with at last 1 million reads). These sequences were uploaded to MG-RAST (58) for annotation. We used Metaxa2 (59) with default settings to extract SSU rRNA gene sequences (bacterial, archaeal, and eukaryotic) in each sample and assigned taxonomy as described above using the Greengenes 13_8 database (56) and the RDP classifier (57). All statistical analyses were done in R studio and all figures were created with ggplot2 (60) unless otherwise noted.

### Assembly, annotation, and characterization of Dormibacteraeota genomes

We assembled two genomes belonging to the candidate phylum Dormibacteraeota (29) from individual metagenomes obtained from Calhoun site 1 (60-70cm) and Shale Hills site 1 (90-100 cm). These two soil samples were selected for deeper sequencing based on the high abundance of the phylum Dormibacteraeota (∼60% of amplicon 16S rRNA gene reads at Calhoun, ∼23% at Shale Hills). This sequencing effort yielded 57.7 million paired-end reads for Calhoun 60-70cm and 65.6 million paired-end reads for Shale Hills 90-100cm.

Genomes were assembled using unpaired reads that had been filtered using sickle (-q 20 -l 50). We used Megahit (61) with the bulk preset to build the assembly, and MaxBin 2.2.1 (62) for binning. We used a script to cycle through MaxBin conditions (-min_contig_length 1100 -1500 and -prob_threshold 0.95 - 0.99) and used checkM (63) to pick the best bins. Bins were then manually curated using a combination of scaffold abundance, tetranucleotide frequency, and GC content. After selecting the highest quality bins from each sample, we ran Metaxa2 on the bins themselves to detect SSU or LSU rRNA genes that could be used to determine taxonomic affiliations. Bin 3 contained one 16S rRNA sequence, which matched at 97.2% sequence identity to a Dormibacteraeota sequence within the Greengenes database (sequence id = 151897). The Bin 3 16S rRNA also matched the amplicon sequence for “OTU1” at 98% identity. This OTU was the most abundant AD3 sequence in our amplicon dataset; its maximum relative abundance was 57% at Calhoun Site 1 50cm and it reached ≥1% relative abundance in 43% of our sites. Bin JG-37 contained only small fragments of the 5S and 23S rRNA genes that were insufficient for taxonomic classification.

To verify that these two bins are affiliated with the Dormibacteraeota candidate phylum, we used the concatenated marker gene phylogeny generated from GTDBTk (26) to compare the placement of our genomes to previously published Dormibacteraeota genomes (29, 30). While GTDBTk could only taxonomically classify our genomes to the bacterial domain (a problem replicated in 45/53 currently available Dormibacteraeota genomes), both genomes clearly fall within the Dormibacteraeota phylum in the tree generated from concatenated marker genes (Figure 4B). This tree was plotted in ITOL (64). Both Dormibacteraeota genomes were submitted to IMG for annotation under the taxon IDs 2756170100 (binJG37, please see note in data availability section) and 2767802471 (bin3). Based on CheckM (63) estimates, both genomes are substantially complete with medium to high contamination (bin JG37: 69.9% complete, 7.7% contamination; bin3: 72.6% complete, 10.7% contamination) See Supplemental Table 1 for additional genome details.

### Phylogenetic tree of CoxL genes

The evolutionary history of the Dormibacteraeota coxL genes was inferred by using the Maximum Likelihood method based on the JTT matrix-based model (65). The tree with the highest log likelihood (-24038) is shown. The percentage of trees in which the associated taxa clustered together is shown below the branches. Initial tree(s) for the heuristic search were obtained automatically by applying Neighbor-Join and BioNJ algorithms to a matrix of pairwise distances estimated using a JTT model, and then selecting the topology with superior log likelihood value. A discrete gamma distribution was used to model evolutionary rate differences among sites (5 categories (+G, parameter = 1.10)). The rate variation model allowed for some sites to be evolutionarily invariable ([+I], 6.75% sites). The resulting tree was drawn to scale, with branch lengths measured in the number of substitutions per site. The analysis involved 67 amino acid sequences. All positions containing gaps and missing data were eliminated. There were a total of 526 positions in the final dataset. Evolutionary analyses were conducted in MEGA7 (66) and the tree was plotted in ITOL (64).

### Calculation of maximum growth rate proxy ΔENC’

Because there are no cultivated members of phylum Dormibacteraeota, we calculated ΔENC’ to estimate potential growth rates as described previously (40, 41). We also calculated ΔENC’ on complete genomes in NCBI that matched amplicon sequences in our dataset with >=99% sequence similarity. We used this set of genomes to represent the bacteria found in the same soil profiles studied here to establish a range for microbial growth rates in soil. We ran ENCprime (40) with default options on both concatenated ribosomal protein sequences and concatenated genome sequences, and calculated ΔENC’ as described in Vieira-Silva (2009).

### Spore selection treatment

We adapted a method previously used in human stool samples (48) to select for spores in a culture independent manner in three soil samples from our study (Calhoun site 1, soils 50-60cm, 60-70cm, and Calhoun site 2, soil 50-60cm). To select for spores, we incubated 0.04g of each sieved soil (as described above) in 70% ethanol for 4 hours under aerobic conditions and constant agitation with the goal of killing vegetative cells. After the incubations, we washed both sets of samples with PBS three times, then applied propidium monoazide (PMA) to the ethanol-treated samples as described previously (49). We used PMA to block the amplification of DNA from cells with compromised membranes, ensuring that only those cells capable of surviving the harsh ethanol treatment would be amplified in subsequent PCRs. We PCR amplified, sequenced, and processed these samples as previously described. We restricted our analysis to the top 1000 most abundant phylotypes to remove rare taxa and used the Wilcoxon test to identify enriched taxa, scoring taxa as “possible spore formers” if they had False Discovery Rate (FDR) corrected p-values greater than 0.05. These taxa are presented in Supplemental Table 3.

### Data availability

Both Dormibacteraeota genomes are publicly available on IMG under taxon IDs 2767802471 (bin3) and 2756170100 (bin JG37). Bin JG37 was refined and resubmitted to IMG (on 01May2019), and may not be updated by the time this manuscript is reviewed. The refined bin JG37 is available on figshare at https://doi.org/10.6084/m9.figshare.8156261.v1, along with a list of contigs removed from the original bin JG37, allowing reviewers to view changes in IMG. The metagenomes these genomes were assembled from all also publicly available on IMG under taxon ID 3300022691 (Calhoun 60cm) and 3300021046 (Shale hills 90cm). The merged, quality filtered, and unassembled shotgun sequences are available under MG-RAST project ID mgp80869. The raw, unmerged 16S amplicon sequences are available on figshare at https://doi.org/10.6084/m9.figshare.4702711.

## Acknowledgements

We want to thank Chelsea Carey, Neal Blair, and Andrew Bissett for their contributions to this research effort and Belinda Ferrari for providing feedback on a previous draft of this manuscript. Funding for this work was provided the U.S. National Science Foundation’s Critical Zone Observatories program with additional funding provided by the NSF EarthCube program (ICER-1541047) and the Macrosystems program (EF-1550920).

## Author Contributions

All authors contributed to this project by collecting/processing samples, characterizing soils, extracting DNA, analyzing data, or some combination thereof. E.A. was primarily responsible for leading this cross-site effort and for coordinating the research activities across all project personnel. T.E.B. led the amplicon and shotgun metagenomic analyses along with the associated bioinformatics and data analyses. The manuscript was written by T.E.B. and N.F. with input from all co-authors.

## Supplementary Information for

### This section includes

Figs. S1 to S4

Tables S1 to S3

**Fig. S1.**
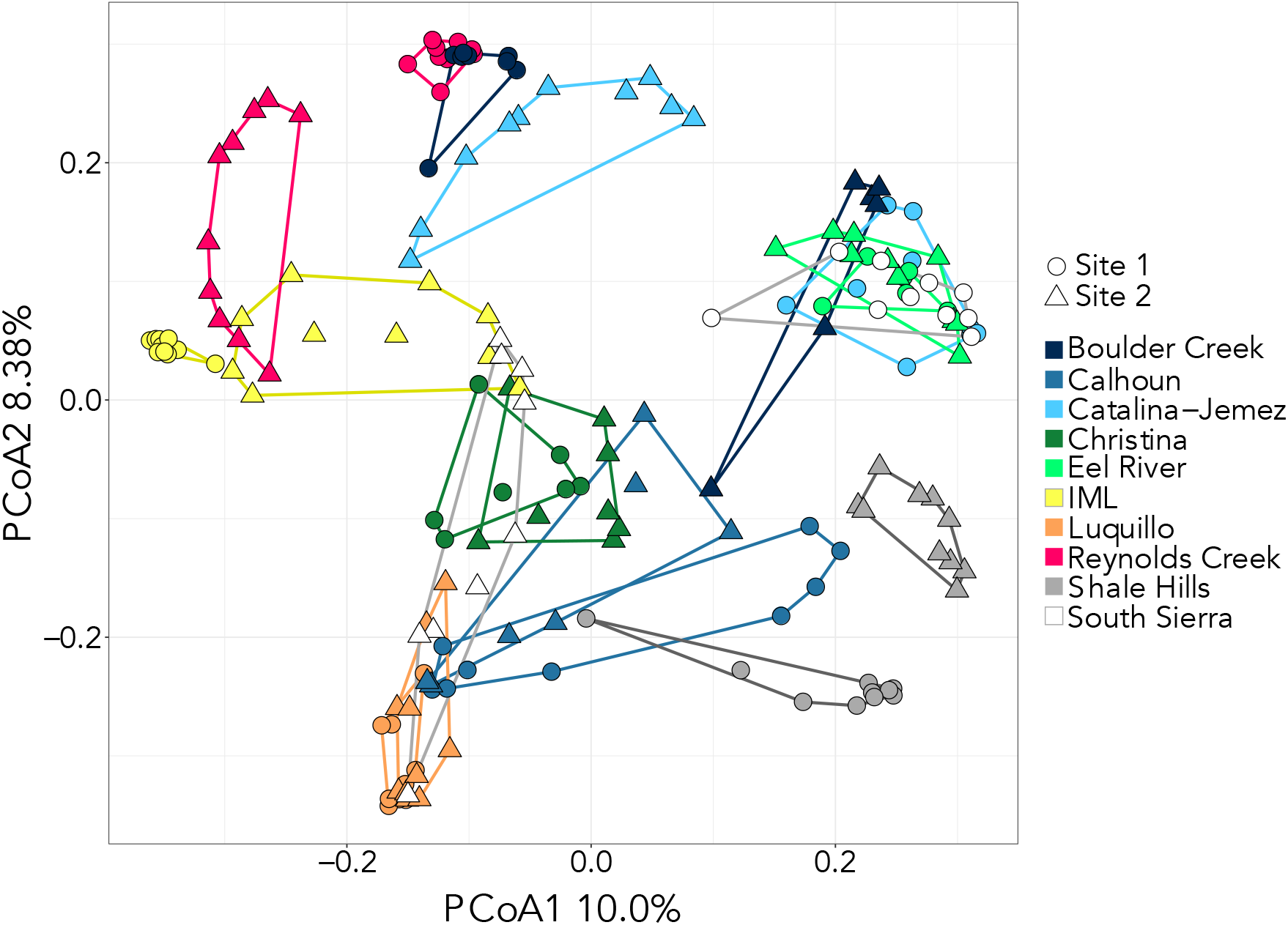
Ordination plot showing differences in overall microbial community composition across the 20 sampled profiles (two per Critical Zone Observatory). The principal coordinates analysis is based on Bray-Curtis dissimilarities calculated from the 16S rRNA gene amplicon data. This ordination plot shows that the differences in communities between profiles are typically larger than the differences in communities across different depths within individual profile, a conclusion supported by the associated PerMANOVA analyses.

**Fig. S2.**
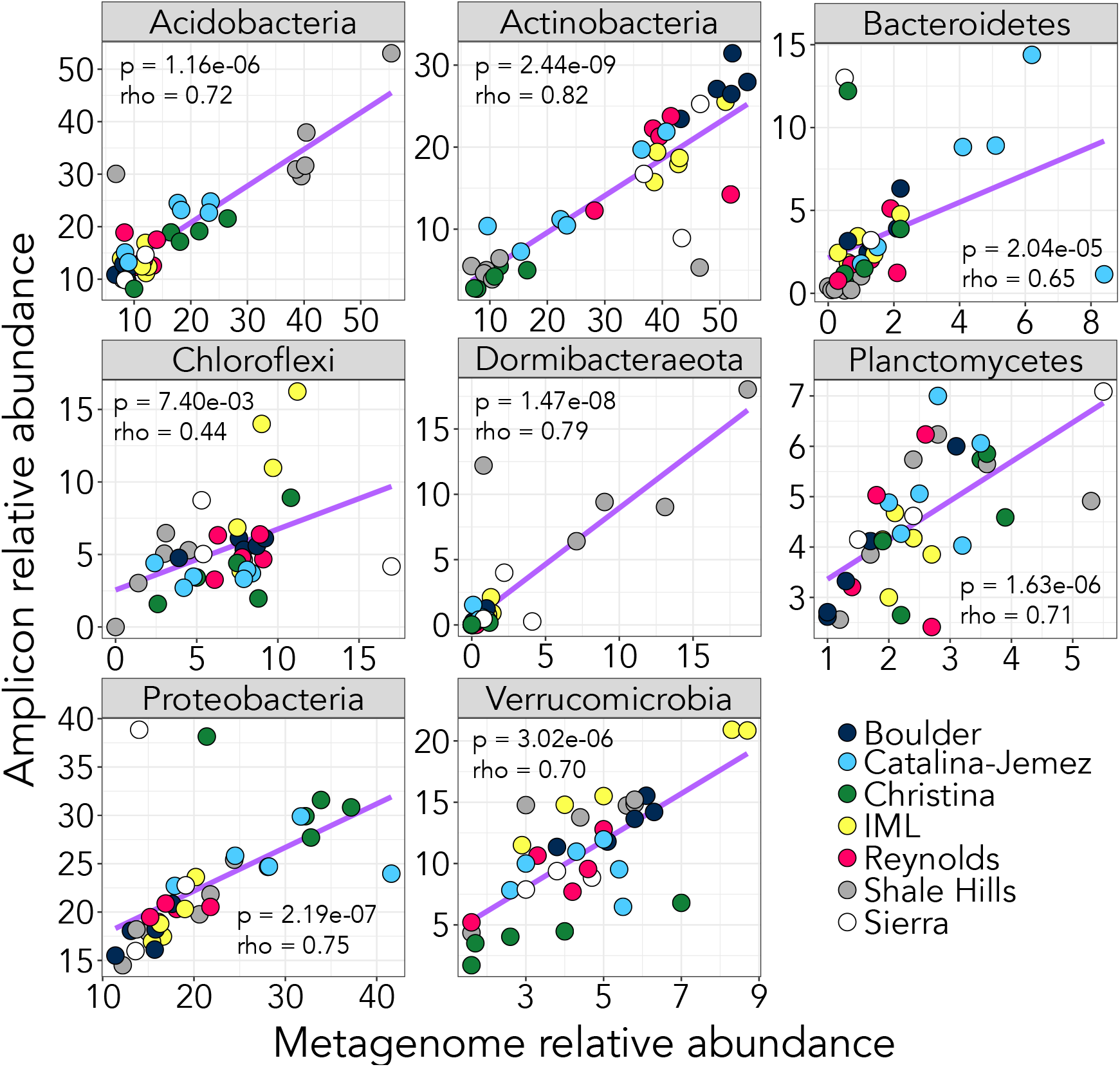
Relative abundances of the eight most abundant bacterial phyla in our dataset are well correlated between 16S rRNA gene amplicon and shotgun metagenomic methods. We used Metaxa2 (59) to search for SSU rRNA gene fragments in our metagenomic data. P-values and rho values indicate Spearman correlations.

**Fig. S3:**
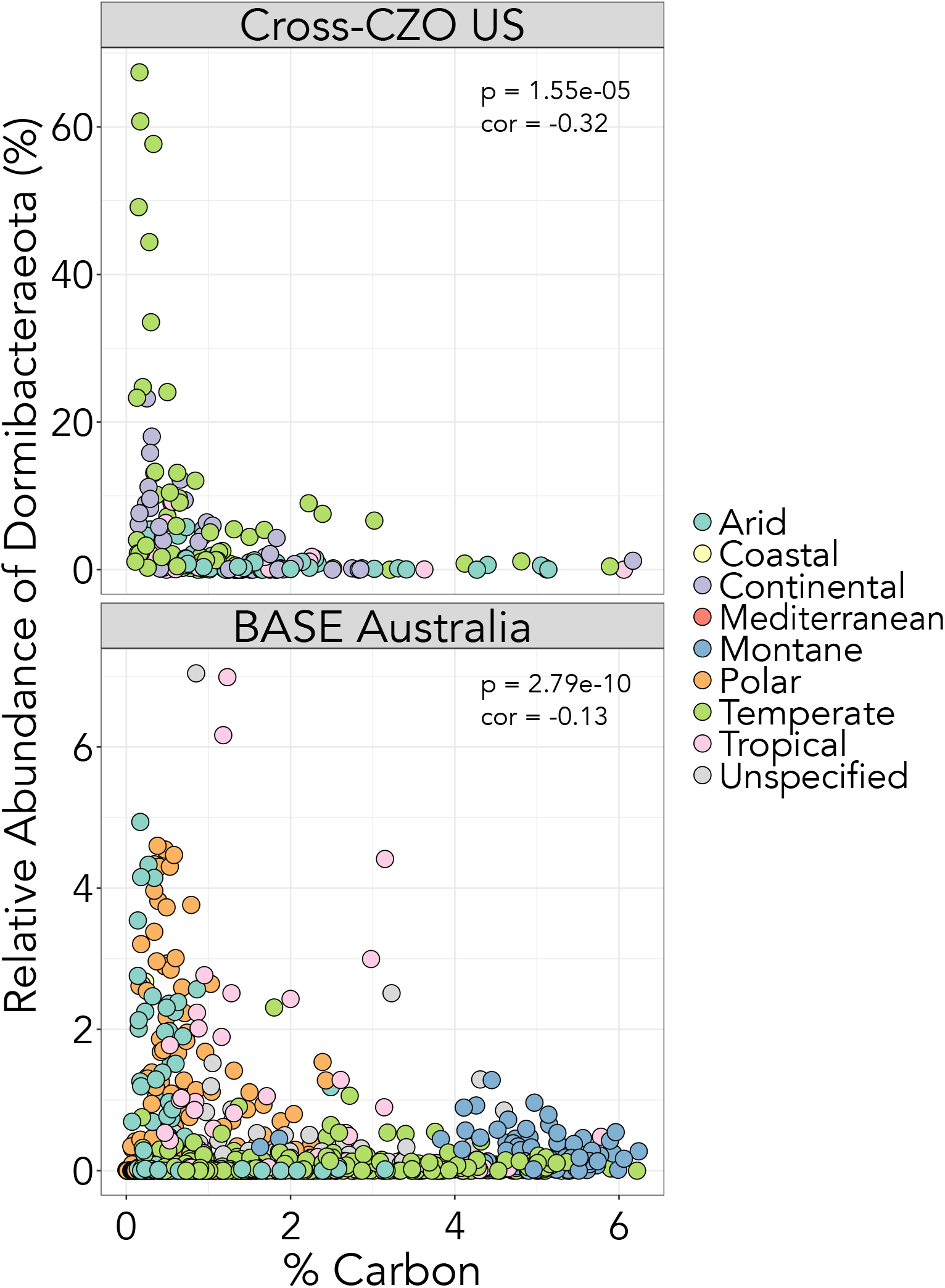
The relative abundance of phylum Dormibacteraeota is negatively correlated with soil carbon concentrations across two independent soil datasets. The top panel features data from this study (20 soil profiles across the U.S., 179 soils in total) while the bottom panel draws from a dataset encompassing 1006 surface soils (0-10 cm depth) collected from across Australia as part the BASE project (Biomes of Australian Soil Environments; (33). P-values and correlation coefficients (cor) are from Pearson correlations.

**Fig. S4:**
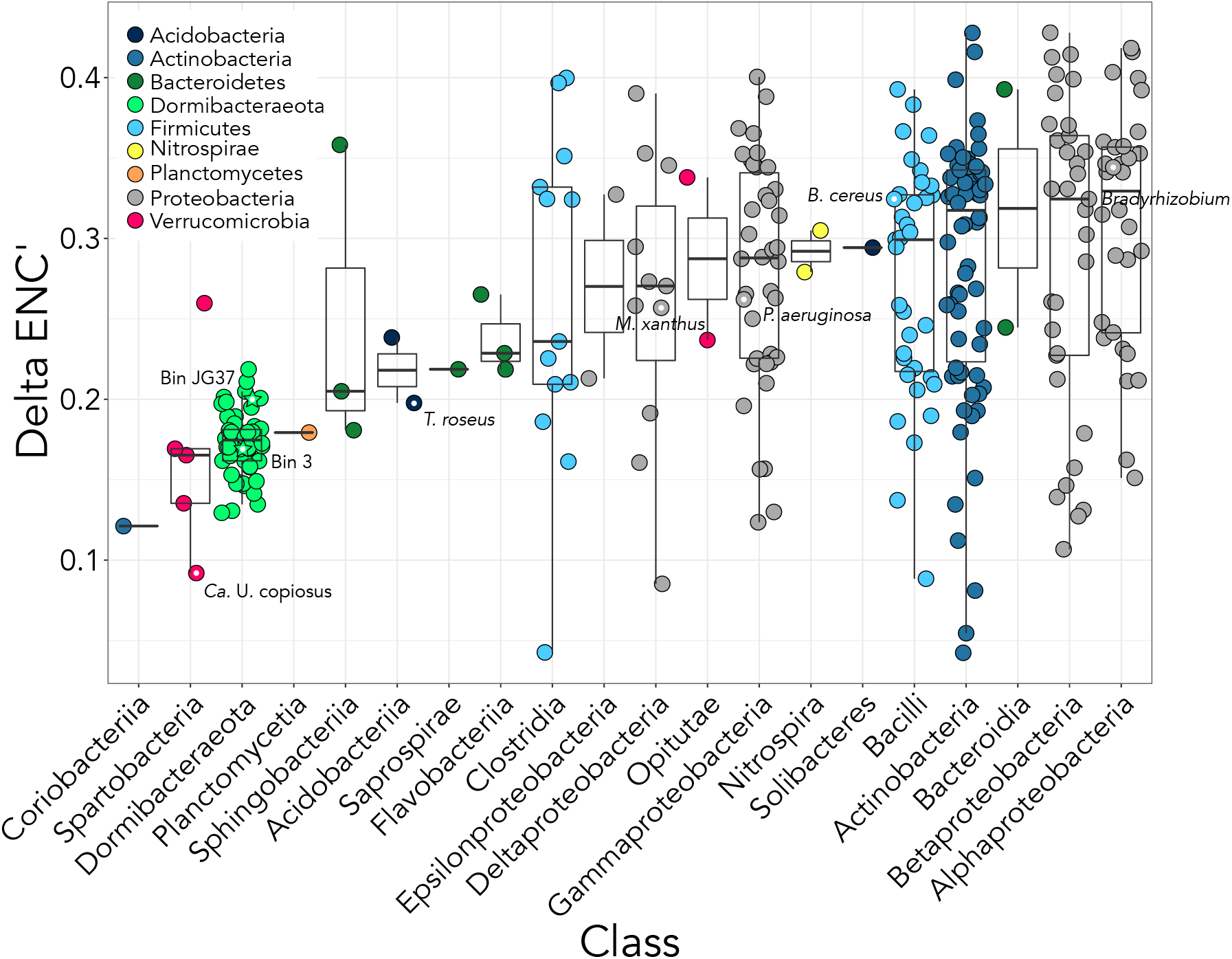
Members of the phylum Dormibacteraeota are predicted to have low maximum potential growth rates based on the growth rate proxy ΔENC’ (a metric of codon usage bias). ΔENC’ ranges from our Dormibacteraeota genomes, the Antarctic Dormibacteraeota genomes (29), the thawing permafrost Dormibacteraeota genomes (30), and a set of genomes matched to our 16S rRNA gene amplicon sequences are shown, arranged by taxonomic affiliation. ΔENC’ is positively correlated with growth rate in bacterial and archaeal genomes (41). All Dormibacteraeota genomes originated from similar carbon-limited environments, which may account for the tight clustering of ΔENC’ values for this phylum compared to the other presented classes. Select genomes are labeled for additional context and indicated with a central white dot.

**Fig. S5:**
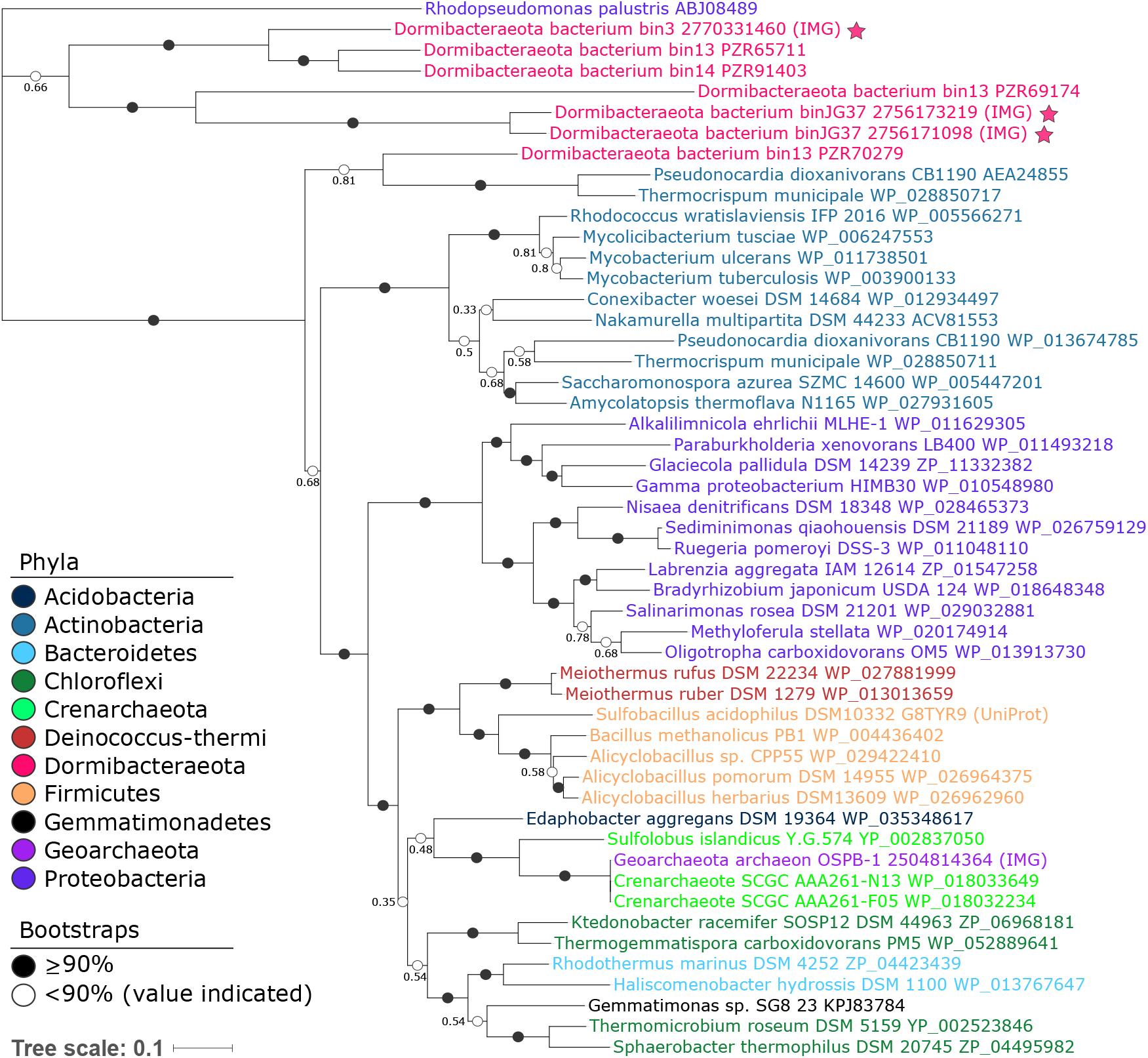
The Dormibacteraeota genomes assembled from cross-CZO soil metagenomes contain form II CO dehydrogenases (coxL). Form II coxL genes may or may not be associated with the ability to oxidize carbon monoxide (42). However, the Antarctic bin13 Dormibacteraeota genome (29) contains a form I coxL sequence most closely related to Actinobacteria, indicating that at least some members of this phylum are likely capable of CO oxidation. Proteins included in this figure are indicated by their Genbank IDs unless otherwise indicated. Proteins from genomes assembled in this study are indicated with stars. Details on the tree construction are included in the Methods text.

**Table S1.**
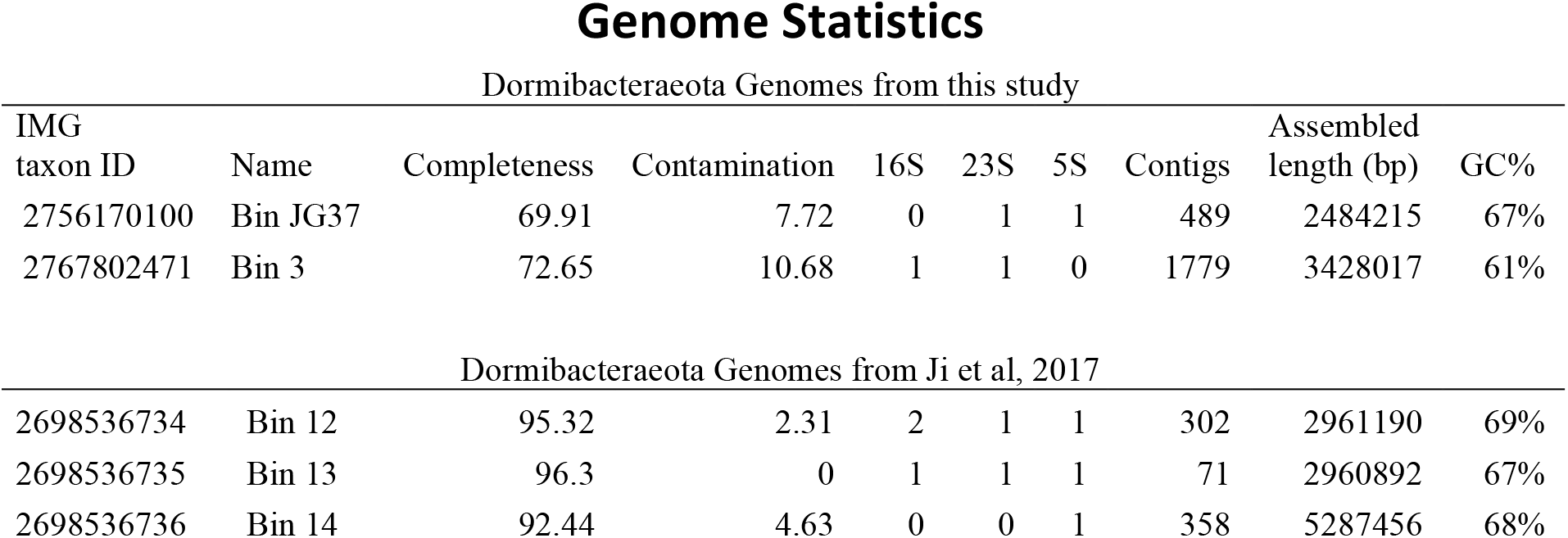
Genome statistics from genomes assembled in this study and Dormibacteraeota genomes assembled in Ji et al, 2017. Bin JG37 was refined and resubmitted to IMG - see data availability for more details.

Dormibacteraeota Genomes from this study
| IMG<br>taxon ID | Name | Completeness | Contamination | 16S | 23S | 5S | Contigs | Assembled<br>length (bp) | GC% |
| --- | --- | --- | --- | --- | --- | --- | --- | --- | --- |
| 2756170100 | Bin JG37 | 69.91 | 7.72 | 0 | 1 | 1 | 489 | 2484215 | 67% |
| 2767802471 | Bin 3 | 72.65 | 10.68 | 1 | 1 | 0 | 1779 | 3428017 | 61% |
| Dormibacteraeota Genomes from Ji et al, 2017 |  |  |  |  |  |  |  |  |  |
| 2698536734 | Bin 12 | 95.32 | 2.31 | 2 | 1 | 1 | 302 | 2961190 | 69% |
| 2698536735 | Bin 13 | 96.3 | 0 | 1 | 1 | 1 | 71 | 2960892 | 67% |
| 2698536736 | Bin 14 | 92.44 | 4.63 | 0 | 0 | 1 | 358 | 5287456 | 68% |

**Table S2.**
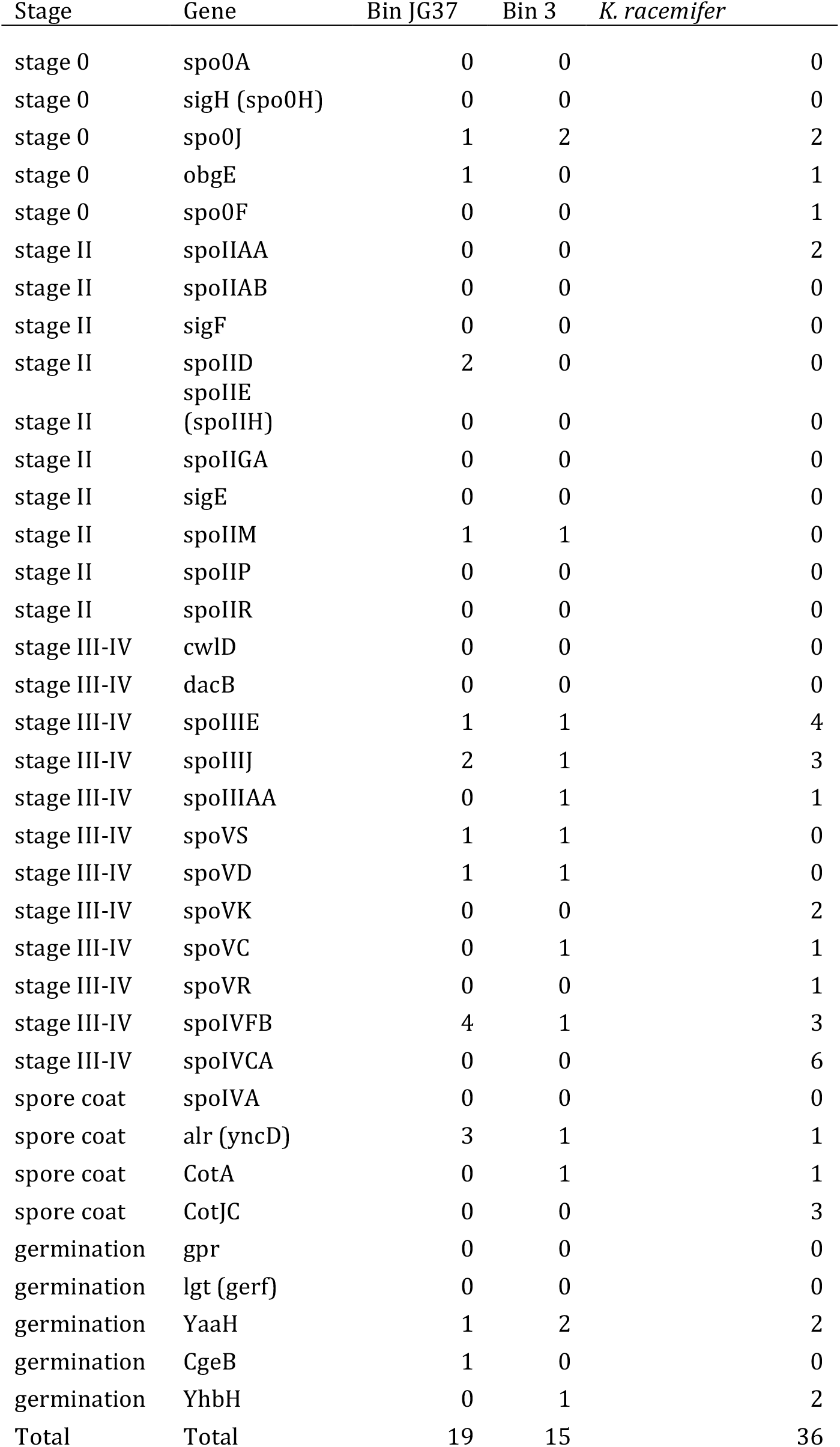
Spore forming genes from Dormibacteraeota genomes and closest spore-forming relative *K. racemifer*.

**Table S3:**
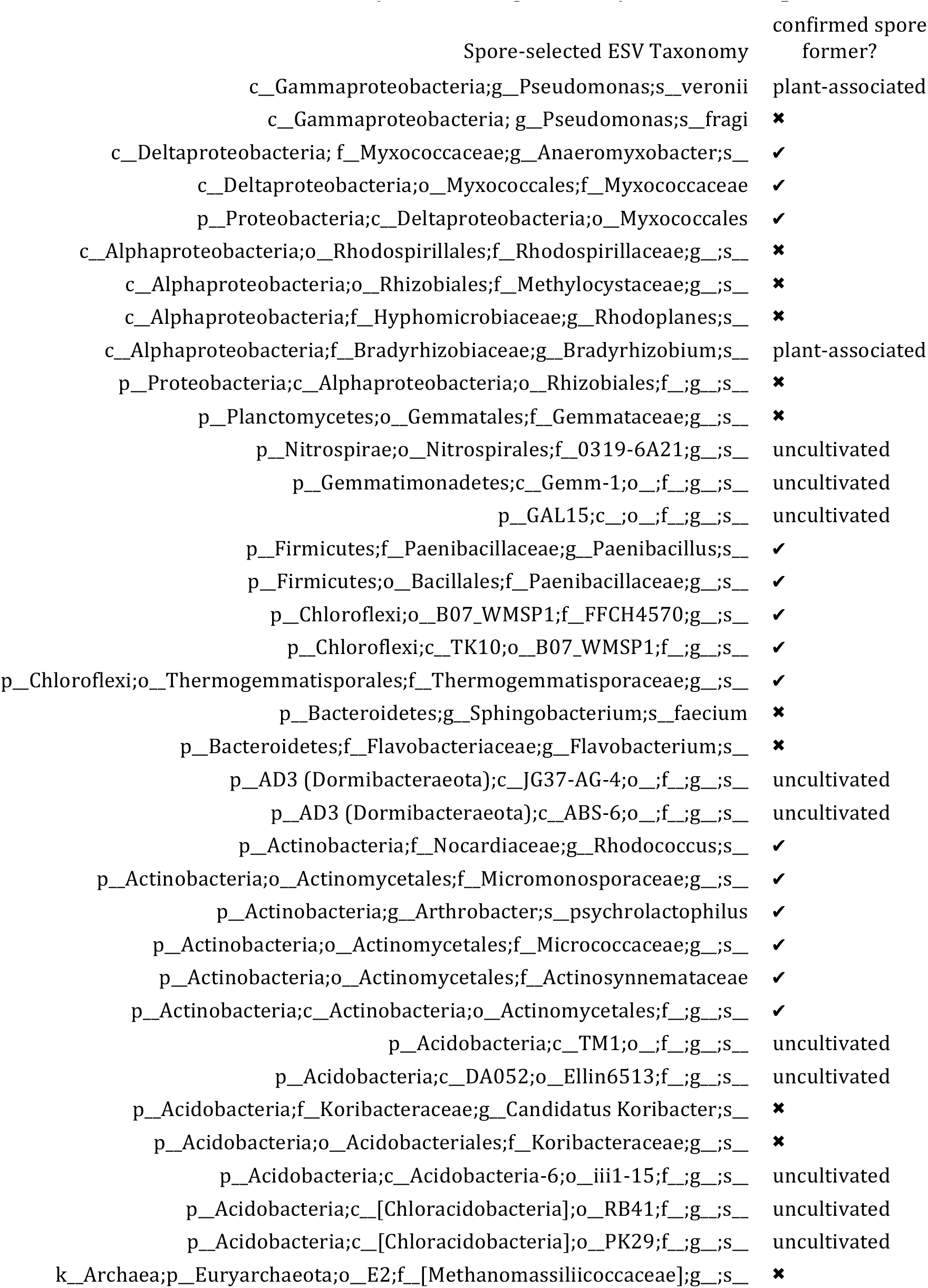
Taxonomy of ESVs significantly enriched in spore selection treatment.

**Additional data table S1 (separate file)**

**Site details - chemical, texture, and metadata for all samples included in study**

**Additional data table S2 (separate file)**

**Phylum level abundances of all bacteria and archaea derived from 16S amplicon data**

## References

1. Jobbágy EG, Jackson RB (2000) The vertical distribution of soil organic carbon and its relation to climate and vegetation. Ecological Applications 10(2):423–436.

2. Balesdent J, et al. (2018) Atmosphere–soil carbon transfer as a function of soil depth. Nature 559:599–602.

3. Schütz K, Kandeler E, Nagel P, Scheu S, Ruess L (2010) Functional microbial community response to nutrient pulses by artificial groundwater recharge practice in surface soils and subsoils. FEMS Microbiol Ecol 72(3):445–455.

4. Eilers KG, Debenport S, Anderson S, Fierer N (2012) Digging deeper to find unique microbial communities: The strong effect of depth on the structure of bacterial and archaeal communities in soil. Soil Biology and Biochemistry 50(C):58–65.

5. Fierer N, Schimel JP, Holden PA (2003) Variations in microbial community composition through two soil depth profiles. Soil Biology and Biochemistry 35:167–176.

6. Blume E, et al. (2002) Surface and subsurface microbial biomass, community structure and metabolic activity as a function of soil depth and season. Applied Soil Ecology 20:171–181.

7. Spohn M, Klaus K, Wanek W, Richter A (2016) Microbial carbon use efficiency and biomass turnover times depending on soil depth - Implications for carbon cycling. Soil Biology and Biochemistry 96(C):74–81.

8. Stone MM, DeForest JL, Plante AF (2014) Changes in extracellular enzyme activity and microbial community structure with soil depth at the Luquillo Critical Zone Observatory. Soil Biology and Biochemistry 75(c):237–247.

9. Kramer C, Gleixner G (2008) Soil organic matter in soil depth profiles: Distinct carbon preferences of microbial groups during carbon transformation. Soil Biology and Biochemistry 40(2):425–433.

10. Banning NC, Maccarone LD, Fisk LM, Murphy DV (2015) Ammonia-oxidising bacteria not archaea dominate nitrification activity in semi-arid agricultural soil. Nature 5(11146):1–8.

11. Oh N-H, Richter DD (2005) Elemental translocation and loss from three highly weathered soil–bedrock profiles in the southeastern United States. Geoderma 126(1-2):5–25.

12. Fimmen RL, Richter DD Jr., Vasudevan D, Williams MA, West LT (2008) Rhizogenic Fe–C redox cycling: a hypothetical biogeochemical mechanism that drives crustal weathering in upland soils. Biogeochemistry 87(2):127–141.

13. Hall SJ, Liptzin D, Buss HL, DeAngelis K, Silver WL (2016) Drivers and patterns of iron redox cycling from surface to bedrock in a deep tropical forest soil: a new conceptual model. Biogeochemistry 130(1):177–190.

14. Schwarz A, et al. (2018) Microbial Degradation of Phenanthrene in Pristine and Contaminated Sandy Soils. Microb Ecol 75:888–902.

15. Will C, et al. (2010) Horizon-specific bacterial community composition of German grassland soils, as revealed by pyrosequencing-based analysis of 16S rRNA genes. Applied and Environmental Microbiology 76(20):6751–6759.

16. Kramer S, Marhan S, Haslwimmer H, Ruess L, Kandeler E (2013) Temporal variation in surface and subsoil abundance and function of the soil microbial community in an arable soil. Soil Biology and Biochemistry 61(C):76–85.

17. Tas N, et al. (2014) Impact of fire on active layer and permafrost microbial communities and metagenomes in an upland Alaskan boreal forest. ISME Journal 8:1904–1919.

18. Fierer N (2017) Embracing the unknown: disentangling the complexities of the soil microbiome. Nat Rev Micro :1–12.

19. Vartoukian SR, Palmer RM, Wade WG (2010) Strategies for culture of “unculturable” bacteria. FEMS Microbiology Letters 309:1–7.

20. Marty C, Houle D, Gagnon C, Courchesne F (2017) The relationships of soil total nitrogen concentrations, pools and C:N ratios with climate, vegetation types and nitrate deposition in temperate and boreal forests of eastern Canada. Catena 152(C):163–172.

21. Lloyd KG, Steen AD, Ladau J, Yin J, Crosby L (2018) Phylogenetically Novel Uncultured Microbial Cells Dominate Earth Microbiomes. mSystems 3(5):1–12.

22. Lin X, Kennedy D, Fredrickson J, Bjornstad B, Konopka A (2011) Vertical stratification of subsurface microbial community composition across geological formations at the Hanford Site. Environmental Microbiology 14(2):414–425.

23. Leff JW, et al. (2015) Consistent responses of soil microbial communities to elevated nutrient inputs in grasslands across the globe. Proc Natl Acad Sci USA 112(35):10967–10972.

24. 130.Fierer N, et al. (2011) Comparative metagenomic, phylogenetic and physiological analyses of soil microbial communities across nitrogen gradients. ISME Journal 6(5):1007–1017.

25. Turner S, et al. (2017) Microbial community dynamics in soil depth profiles over 120,000 years of ecosystem development. Front Microbiol 8:1– 17.

26. Parks DH, et al. (2018) A standardized bacterial taxonomy based on genome phylogeny substantially revises the tree of life. Nat Biotechnol 36(10):996–1004.

27. Eloe-Fadrosh EA, Ivanova NN, Woyke T, Kyrpides NC (2016) Metagenomics uncovers gaps in amplicon-based detection of microbial diversity. Nature Microbiology 1(4):1–4.

28. Zhou J, et al. (2003) Bacterial phylogenetic diversity and a novel candidate division of two humid region, sandy surface soils. Soil Biology and Biochemistry 35(7):915–924.

29. Ji M, et al. (2017) Atmospheric trace gases support primary production in Antarctic desert surface soil. Nature 552(7685):400–403.

30. Woodcroft BJ, et al. (2018) Genome-centric view of carbon processing in thawing permafrost. Nature 560:49–54.

31. Kim HM, et al. (2014) Bacterial community structure and soil properties of a subarctic tundra soil in Council, Alaska. FEMS Microbiol Ecol 89(2):465–475.

32. Billings SA, et al. (2018) Loss of deep roots limits biogenic agents of soil development that are only partially restored by decades of forest regeneration. Elementa 6(34):1–19.

33. Bissett A, et al. (2016) Introducing BASE: the Biomes of Australian Soil Environments soil microbial diversity database. GigaScience 5(21):1– 11.

34. Bowers RM, et al. (2017) Minimum information about a single amplified genome (MISAG) and a metagenome-assembled genome (MIMAG) of bacteria and archaea. Nat Biotechnol 35(8):725–731.

35. Konstantinidis KT, Tiedje JM (2005) Towards a genome-based taxonomy for prokaryotes. Journal of Bacteriology 187(18):6258–6264.

36. Fung T, Kwong N, van der Zwan T, Wu M (2013) Residual Glycogen Metabolism in *Escherichia coli* is Specific to the Limiting Macronutrient and Varies During Stationary Phase. Journal of Experimental Microbiology and Immunology JEMI 17:83–87.

37. Kandror O, DeLeon A, Goldberg AL (2002) Trehalose synthesis is induced upon exposure of Escherichia coli to cold and is essential for viability at low temperatures. Proc Natl Acad Sci USA 99(15):1–6.

38. Wilson WA, et al. (2010) Regulation of glycogen metabolism in yeast and bacteria. FEMS Microbiol Rev 34(6):952–985.

39. Roller BRK, Stoddard SF, Schmidt TM (2016) Exploiting rRNA operon copy number to investigate bacterial reproductive strategies. Nature Microbiology 1:1–7.

40. Novembre JA (2002) Accounting for Background Nucleotide Composition When Measuring Codon Usage Bias. Molecular Biology and Evolution 19(8):1390–1394.

41. Vieira-Silva S, Rocha E (2009) The Systemic Imprint of Growth and Its Uses in Ecological (Meta)Genomics. PLOS Genetics 6(1):1–15.

42. King GM, Weber CF (2007) Distribution, diversity and ecology of aerobic CO-oxidizing bacteria. Nat Rev Micro 5(2):107–118.

43. Lorite MJ, Tachil J, Sanjuán J, Meyer O, Bedmar EJ (2000) Carbon Monoxide Dehydrogenase Activity in Bradyrhizobium japonicum. Applied and Environmental Microbiology 66(5):1871–1876.

44. Hille R (2005) Molybdenum-containing hydroxylases. Archives of Biochemistry and Biophysics 433(1):107–116.

45. Cordero PRF, et al. (2019) Carbon monoxide dehydrogenases enhance bacterial survival by oxidising atmospheric CO. bioRxiv.

46. Fujita M, Losick R (2005) Evidence that entry into sporulation in Bacillus subtilis is governed by a gradual increase in the level and activity of the master regulator Spo0A. Genes Development 19:2236–2244.

47. Cavaletti L, et al. (2006) New lineage of filamentous, spore-forming, gram-positive bacteria from soil. Applied and Environmental Microbiology 72(6):4360–4369.

48. Browne HP, et al. (2016) Culturing of “unculturable” human microbiota reveals novel taxa and extensive sporulation. Nature 533(7604):543– 546.

49. Carini P, et al. (2016) Relic DNA is abundant in soil and obscures estimates of soil microbial diversity. Nature Microbiology 2(0):1–6.

50. Shimkets LJ (1999) Intercellular signaling during fruiting-body development of *Myxococcus xanthus*. Annu Rev Microbiol 53:1–26.

51. Yabe S, Aiba Y, Sakai Y, Hazaka M, Yokota A (2011) Thermogemmatispora onikobensis gen. nov., sp. nov. and Thermogemmatispora foliorum sp. nov., isolated from fallen leaves on geothermal soils, and description of Thermogemmatisporaceae fam. nov. and Thermogemmatisporales ord. nov. within the class Ktedonobacteria. International Journal of Systemic and Evolutionary Microbiology 61(4):903–910.

52. Luther MK, Bilida S, Mermel LA, LaPlante KL (2015) Ethanol and Isopropyl Alcohol Exposure Increases Biofilm Formation in *Staphylococcus aureus* and *Staphylococcus epidermidis*. Infectious Diseases and Therapy 4(2):219–226.

53. Zinniel DK, et al. (2002) Isolation and characterization of endophytic colonizing bacteria from agronomic crops and prairie plants. Applied and Environmental Microbiology 68(5):2198–2208.

54. Liptzin D, Silver WL (2009) Effects of carbon additions on iron reduction and phosphorus availability in a humid tropical forest soil. Soil Biology and Biochemistry 41(8):1696–1702.

55. Edgar RC (2016) UNOISE2: improved error-correction for Illumina 16S and ITS amplicon sequencing. bioRxiv:1–21.

56. DeSantis TZ, et al. (2006) Greengenes, a Chimera-Checked 16S rRNA Gene Database and Workbench Compatible with ARB. Applied and Environmental Microbiology 72(7):5069–5072.

57. Cole JR, et al. (2013) Ribosomal Database Project: data and tools for high throughput rRNA analysis. Nucleic Acids Research 42(D1):D633–D642.

58. Meyer F, et al. (2008) The metagenomics RAST server – a public resource for the automatic phylogenetic and functional analysis of metagenomes. BMC Bioinformatics 9(1):386–8.

59. Bengtsson-Palme J, et al. (2015) metaxa2: improved identification and taxonomic classification of small and large subunit rRNA in metagenomic data. Mol Ecol Resour 15(6):1403–1414.

60. Wickham H (2009) ggplot2: Elegant Graphics for Data Analysis (Use R) (Springer).

61. Li D, Liu C-M, Luo R, Sadakane K, Lam T-W (2015) MEGAHIT: an ultra-fast single-node solution for large and complex metagenomics assembly via succinct de Bruijn graph. Bioinformatics 31(10):1674–1676.

62. Wu Y-W, Simmons BA, Singer SW (2016) MaxBin 2.0: an automated binning algorithm to recover genomes from multiple metagenomic datasets. Bioinformatics 32(4):605–607.

63. Parks DH, Imelfort M, Skennerton CT, Hugenholtz P, Tyson GW (2015) CheckM: assessing the quality of microbial genomes recovered from isolates, single cells, and metagenomes. Genome Res 25(7):1043–1055.

64. Letunic I, Bork P (2016) Interactive tree of life (iTOL) v3: an online tool for the display and annotation of phylogenetic and other trees. Nucleic Acids Research 44(W1):W242–W245.

65. Jones DT, Taylor WR, Thornton JM (1992) The rapid generation of mutation data matrices from protein sequences. Computer Application in the Biosciences 8:275–282.

66. Kumar S, Stecher G, Tamura K (2016) MEGA7: Molecular Evolutionary Genetics Analysis Version 7.0 for Bigger Datasets. Molecular Biology and Evolution 33(7):1870–1874.

